# CGCom: a framework for inferring Cell-cell Communication based on Graph Neural Network

**DOI:** 10.1101/2023.11.10.566642

**Authors:** Honglin Wang, Chenyu Zhang, Seung-Hyun Hong, Peter Maye, David Rowe, Dong-Guk Shin

**Affiliations:** Computer Science and Engineering Department, University of Connecticut, Storrs, CT 06269, USA; Department of Reconstructive Sciences, Univ. of Connecticut Health Center, Farmington, USA; Center for Regenerative Medicine and Skeletal Development, Univ. of Connecticut Health Center, Farmington, USA

**Keywords:** Graph neural network, Spatial transcriptomics, Cell-cell Communication

## Abstract

Cell-cell communication is crucial in maintaining cellular homeostasis, cell survival and various regulatory relationships among interacting cells. Thanks to recent advances of spatial transcriptomics technologies, we can now explore if and how cells’ proximal information available from spatial transcriptomics datasets can be used to infer cell-cell communication. Here we present a cell-cell communication inference framework, called CGCom, which uses a graph neural network (GNN) to learn communication patterns among interacting cells by combining single-cell spatial transcriptomic datasets with publicly available ligand-receptor information and the molecular regulatory information down-stream of the ligand-receptor signaling. To evaluate the performance of CGCom, we applied it to mouse embryo seqFISH datasets. Our results demonstrate that CGCom can not only accurately infer cell communication between individual cell pairs but also generalize its learning to predict communication between different cell types. We compared the performance of CGCom with two existing methods, CellChat and CellPhoneDB, and our comparative study revealed both common and unique communication patterns from the three approaches.

Commonly found communication patterns include three sets of ligand-receptor communication relationships, one between surface ectoderm cells and spinal cord cells, one between gut tube cells and endothelium, and one between neural crest and endothelium, all of which have already been reported in the literature thus offering credibility of all three methods. However, we hypothesize that CGCom is superior in reducing false positives thanks to its use of cell proximal information and its learning between specific cell pairs rather than between cell types. CGCom is a GNN-based solution that can take advantage of spatially resolved single-cell transcriptomic data in predicting cell-cell communication with a higher accuracy.

## I Introduction

Multicellular organisms rely on the precise coordination of cellular activities, which hinges upon effective intercellular communication among the various cell types and tissues within the organism [1]–[3]. Thus studying cellular functions increasingly requires how cells communicate with each other given a temporal and spatial context. It is known that signaling events are frequently orchestrated through diverse protein interactions such as ligand-receptor, receptor-receptor, and extracellular matrix-receptor interactions. In addition, the recipient cells, in response, initiate downstream signaling via their respective receptors, typically culminating in the modulation of transcription factor activity and subsequent changes in gene expression.

Historically, communication could only be studied via *in vitro* experiments consisting of one or two cell types and a selected number of genes. Thanks to the advances in single-cell technologies and spatial transcriptomics, which measures gene expression at a single cell resolution, scientists can now attempt to identify cell-cell communication as well as cell type communication. Currently there are many computational tools that can infer cell communication from single-cell transcriptomics. CellChat[4] computes cell type communication probabilities using the law of mass action and considering the geometric means of ligand and receptor expressions weighted by their agonists/antagonists. CellPhoneDB[5] generates a list of most statistically significant ligand-receptor interactions by generating null distribution by permuting cell cluster labels. MISTY[6] constructs a gene communication signaling network within and between cell clusters. SpaOTsc[7] and COMMOT [8] generate cell communication matrix for a given signaling pathway by solving an optimal transport problem. Unfortunately, these methods suffer from two main drawbacks: (i) These methods mostly focus on inferring cell communication at the cell group level as opposed to the single cell level. (ii) The existing methods do not adequately exploit the spatial information among cells and prior knowledge about interacting cells, which are crucial for inferring cell communication because cells are unlikely communicate with each other if they are far away from each other and certain cell types of cells tend to have a high or low probability to communicate with one another. This limitation often introduces false-positive communication patterns.

With the continuous advancement of neural networks, the emerging field of graph neural networks (GNNs) has fundamentally transformed our capacity to comprehend relationships among nodes within graphs by harnessing the power of information propagation from neighboring nodes. We see an opportunity to use this advanced form of neural network to infer cellular communication. Specifically, we hypothesize that the physical locations of cells can be modelled into interconnected graph structures with nodes and edges pairing them and the signal transmission between neighboring cells can modelled into strength discerning the degree of cell-cell interaction. Considering that cells of a specific type display distinct gene expression patterns [9], we also postulate that, to a certain extent, leveraging information on ligand and receptor expressions can also serve as a means to identify cell types and cells’ communication likelihood. For example, a cell must exhibit a specific ligand or receptor expression pattern to be classified as a particular cell type. GNNs are capable of updating cell ligand/receptor expression through interactions with nearby cells. By training the model for cell type classification based on ligand/receptor expression values, we can interpret the relationship coefficients between nodes as communication coefficients when deciphering cell-to-cell communication.

Building on this motivation, we present a deep learning framework called CGCom (Cell GNN-based Communication) designed to model cell-to-cell relationships and the intricate communication patterns that underlie them. The framework takes as input a series directed sub-graph generated from cell physical locations, combined with ligand expression values, and utilizes cell type information as the training objective. The paired cell communication coefficient is computed from the attention scores in the well-trained GAT graph classifier. CGCom then introduces a heuristic computational algorithm to quantify communication between neighboring cells through various ligand-receptor pairs. Our framework demonstrates promising performance in cell classification when evaluated on three mouse embryo seqFISH datasets, using “best practice” cell labels from the same tissue, and it outperforms multilayer perceptron (MLP) baseline. To illustrate the power of CGCom in inferring cell communication, we employ the attention scores from GAT classifier to infer cell communication on the same datasets, revealing common communication patterns between the three datasets. We posit that this GNN-based framework is both powerful and flexible, offering significant potential in predicting cell types and inferring cell-to-cell communication.

## II Methods

### A. Overview of CGCom

An overview of the CGCom workflow is given in Fig. 1. This framework consists of three key computational components: data preprocessing modules, the Graph Attention Network (GAT) graph classifier and the cell communication inference module. In the preprocessing module, the CGCom takes the gene expression value, cell label and cell location to build the dataset for the classifier including the training graph and training node feature values. The GAT graph classifier performs deep learning training to acquire cell type information, leveraging a combination of subgraphs generated from cell spatial locations and preprocessed gene expression values for the cells. Subsequently, the cell communication inference tool utilizes the attention scores from the GAT classifier and incorporates gene expression data to infer communication score between pairs of cells.

**Fig. 1.**
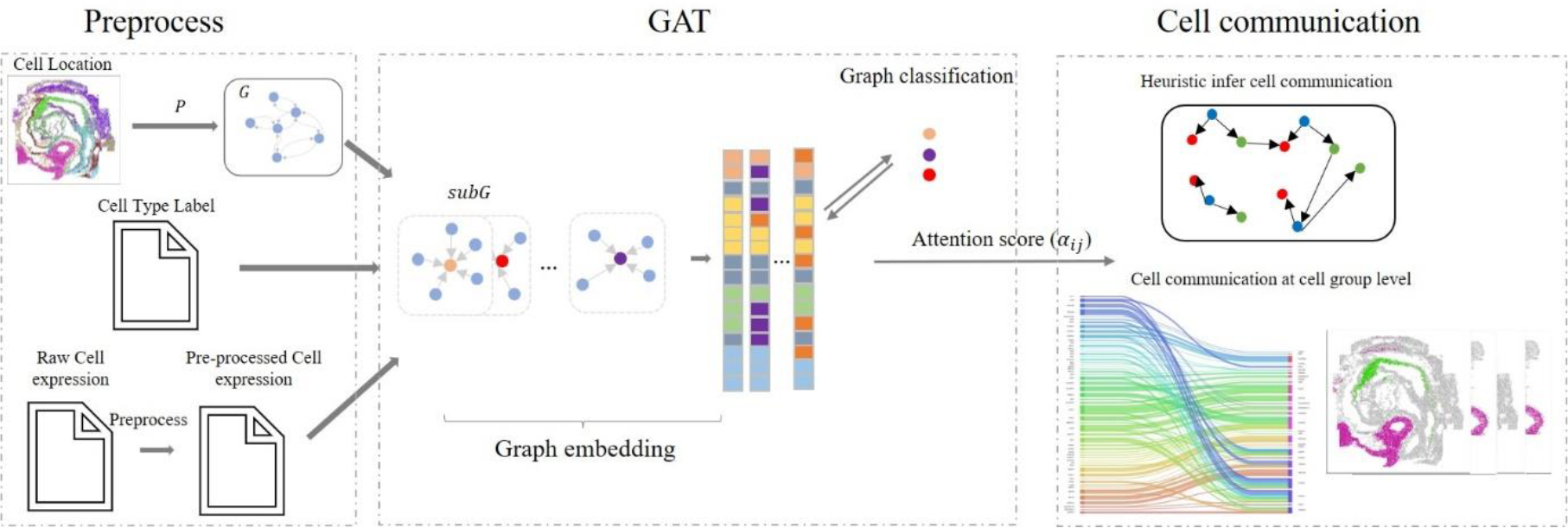
CGCom takes the gene expression matrix, cell type and cell label as the input. The GAT learns the ligand expression patterns of different cell types in a semi-supervised model. The framework then extract the attention score in each graph embedding layer in the GAT from the trained model and infer the communication using a heuristic rule. The communication results finally analyzed at both cell level and cell group level.

### B Curate ligand-receptor-downstream database

Iintercellular responses are initiated with the binding of a ligand to its corresponding receptor, activating specific cell signaling pathways. The foundation for Cell-Cell Communication (CCC) inference lies in constructing a database of ligand-receptor pairs. CGCom uses this database comprised of two parts: 1) ligand and receptor pairs imported from CellChat and CellPhoneDB via direct downloaded from their websites, and 2) receptor and downstream route pairs extracted from the KEGG pathway. The KEGG pathway driven receptor and downstream route pairs are constructed by utilizing a random walk algorithm that begins at the receptor and ends at any encountered transcription factor of the pathway. This data collection process produces a database containing 5842 ligand-receptor-downstream route triplets, including 774 unique ligands, 708 unique receptors, and 308 unique downstream routes.

### C Dataset preprocessing

The dataset preprocessing consists of two primary steps: gene expression preprocessing and cell graph producing. During the gene expression preprocessing phase, the raw scRNA-seq data is filtered and normalized and subject to the quality control process (QC) using Scanpy [10]. The QC includes dealing with gene expression dropouts and mitigating the influence of extreme values. To remedy the high dropout rate inherent in scRNA-seq expression data and to mitigate the influence of extreme values, we use log transformed gene expression of given gene *g* in cell *c* (*v*′_*gc*_), after the original expression value (*v*_*gc*_) is normalized with the non-zero gene-wise median value (*med*(*V*_*g*_=}*v*_*g*1_, …, *v*_*gc*_ (*v*_*gc*_ > 0)})) (Eq. 1). Following this step, the downstream route scores for each cell are computed by applying rPAC [11] to the log-transformed expression values. The pathway scoring system rPAC quantifies the downstream routes by evaluating the consistency between the actual gene regulatory pattern and the expected regulatory pattern for all genes within the route. For a cell *c*, the score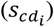 of the downstream route (*d*_*i*_) using all transformed gene expression values 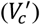 is expressed in Eq. 2.

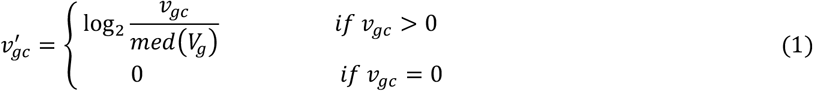

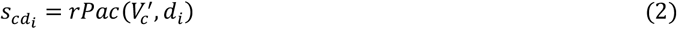

For the cell graph producing, CGCom uses cell locations to construct a directed graph with a hyperparameter *P* designed to limit the maximum communication distance. For the 2-D location of a given cell *c*_*i*_(*x*_*i*_, *y*_*i*_), the graph builder places a node *n*_*i*_ at the corresponding location (*x*_*i*_, *y*_*i*_) within the graph structure (*G*(*N, E*)). Once all nodes are incorporated into the graph, the Euclidean distances *d*_*ij*_ are calculated for each pair of nodes (*i* and *j*) using Eq. 3. We assume that cell communication is restricted to a specific distance range threshold, say *P*. The objective is to introduce directed edges between a pair of nodes in G, if the distance between the nodes is less than the predefined threshold *P*. This cell graph builder uses Algorithm (1) to add directed edges to the graph.

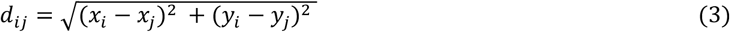

### D The architecture of GAT and the training preprocess

The Graph Attention Network (GAT) [12] is a type of neural network designed for tasks involving graph-structured data, such as graph classification, node classification, and link prediction. GAT uses attention mechanisms to weigh the importance of neighboring nodes when aggregating information from the graph. We compute attention scores of each node in the graph for its neighboring nodes, and then combines their features using these scores. The GAT mechanism can be expressed by Eq. 4 where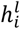 represents the feature vector of node *n*_*i*_ at the *l*th graph embedding layer, *N*(*i*) represents the neighbors of node *i, W*^*l*^ is a weight matrix used for transforming the features of the neighbors, *σ* is an activation function (ReLU or Leaky ReLU) applied element-wise to the aggregated information, and 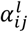represents the attention score for node *i*’s neighbor *j*. These attention scores are computed based on the features of both nodes *i* and *j* and are normalized across all neighbors *j* using softmax function so that the attention score can be easily comparable as shown in Eq. 5. Here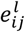 is a learnable parameter associated with the pair of nodes *i* and *j*, and is computed as a function of their features (Eq. 6) where *a*^*l*^ is a learnable weight vector that determines the importance of different aspects of the node features. To simplify the computation step, we use choose ReLU in this equation.

#### Algorithm 1.

Algorithm for adding edge to graph

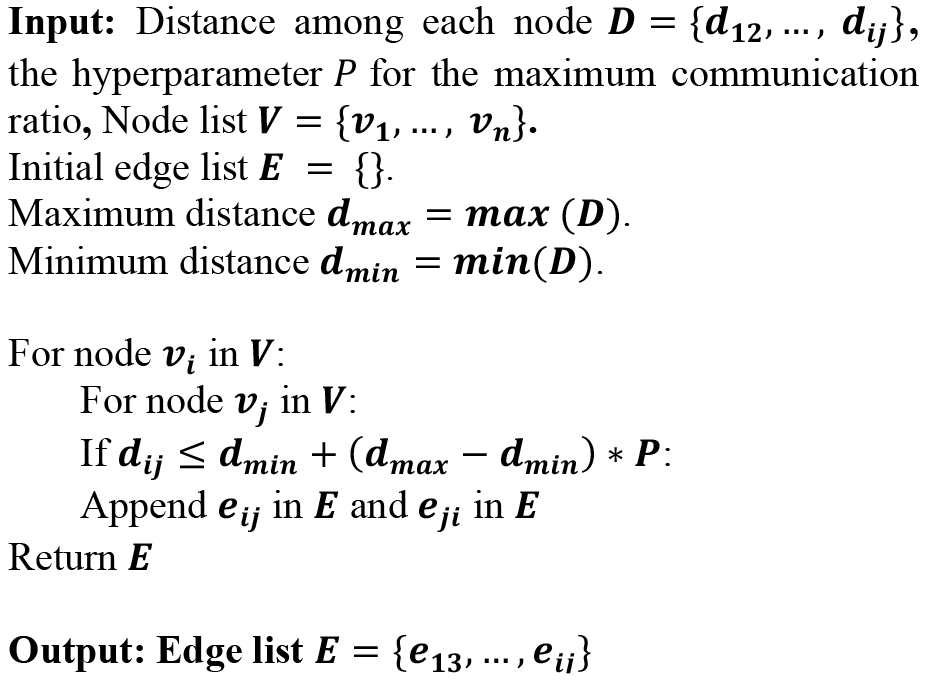

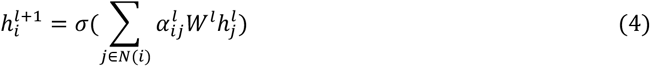

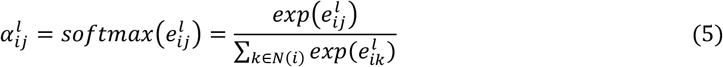

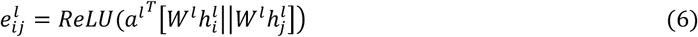

Within this framework, GAT is trained for a graph classification problem. Consequently, the GAT in CGCom is comprised of two components, as illustrated in Fig. 2: the graph embedding part and the graph classification part. The graph embedding part includes three layers, where each layer has the same number of nodes as the input layer. The output of the graph embedding part serves as the input for the graph classification part, which, in turn, includes an output layer with the number of nodes corresponding to the different cell type classes. In a practical situation, it is hard for humans to identify cell type for all cells given an microscopic image, but the gene expression values can be easily obtained using variant sequencing algorithms. Thus, the classifier is trained in a semi-supervised learning mode, using only a few human-labeled data. Since the GAT is used for the graph classification, its dataset is further generated from the two preprocessed data in the preprocessing step. For each node *n*_*i*_ in graph *G*(*N, E*), a subgraph *subG*_*i*_(*N*(*i*), *E*(*i*)) is built with the nodes that has a link with node *n*_*i*_ (*N*(*i*)) and the edge that connecting them (*E*(*i*)). These nodes are also known as the neighbor of node *n*_*i*_ . To stimulated how the expression value of a ligand will impact the cell type classification step, each node in *subG*_*i*_ will have its feature vector (ℎ_*i*_) that only contains all ligands expression values (774 features). The default ratio of training set is 1% and the ratio of validation set is 10%. We use the training set with Adam optimizer and train the model [13] and adjust the learning rate and other hyperparameters with the validation set. To prevent the overfitting, an early stop mechanism is introduced to the training process when the validation loss starts increasing and the validation accuracy starts decreasing.

**Fig. 2.**
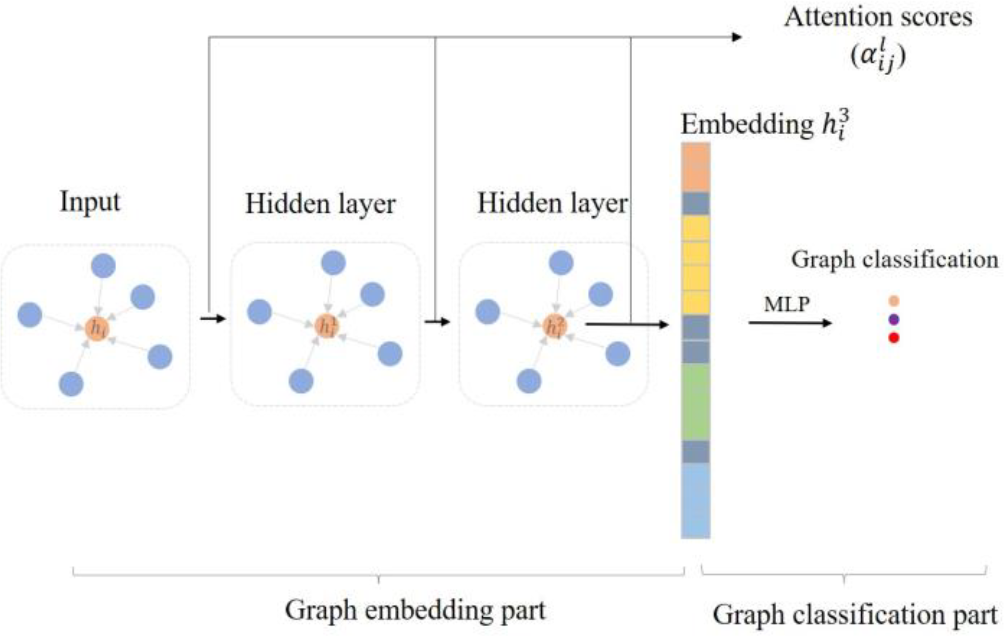
The graph classifier has two including graph embedding part and graph classification part. The graph embedding part embed the neighbor nodes information in the graph into a vector using attention mechanism. Then the graph classification part accepts the embedded vectors and predict the graph type.

### E. The inference of cell communication

After the training iteration stops, we infer the communication with a heuristic rule by intaking the communication coefficient, the ligand and receptor expression value, and corresponding downstream routes in each cell. For each two linked cells *c*_*i*_ and *c*_*j*_, their communication coefficient (*ce*_*ij*_) is computed from the average attention score from three graph embedding layer (Eq. 7). Intuitively both expression values of a ligand of the sender cell and a receptor in the receiver cell should be high, and both expression values of a receptor in sender cell and a ligand in the receiver cell should be low if one were to infer paracrine cell communication. We compute the communication score from *c*_*i*_ to *c*_*j*_ through ligand and receptor pair (*lr*) using Eq. 8 where ligand and receptor pair (*lr*) expression value in both cells *i* and *j* are denoted by (*l*_*i*_, *r*_*i*_) and (*l*_*j*_, *r*_*j*_), respectively, and the downstream route (*d*_*i*_) score is denoted by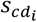 . Below *LRE*(*∎*) used in Eq. 8 is the function computing the ligand and receptor communication strength as shown in Eq. 9. To scaling down the communication score and prevent all the values from becoming close to zero, we use log1p transformed values instead of using the raw value.

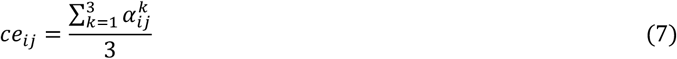

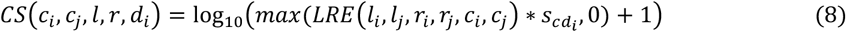

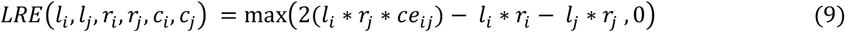

To generalize specific cell-cell communications to cell group-level interactions, we aggregate all the communication scores from group *A* to group *B* by grouping feasible ligand and receptor pairs (*lr*). We employ a permutation test to determine the statistical significance of the inferred cell group communication. The permutation test involves random permutations of cell labels followed by the recalculation of the average group communication score between *A* and *B* via the communication ligand *l* and receptor pair *r* (Eq. 10). Here, *CS*(*A, B, l, r, d*_*i*_) represents the average communication score for cells in group α communicating with cells in group β through a particular ligand and receptor pair (*lr*) (Eq. 11). *CS*′(*A, B, l, r, d*_*i*_) corresponds to the average communication score after the cell labels have been randomly permuted. We repeat this process for a total of *Q* random samples, with *Q* default set to 100 and the default p-value threshold for the statistically significant communication is set to 0.05.

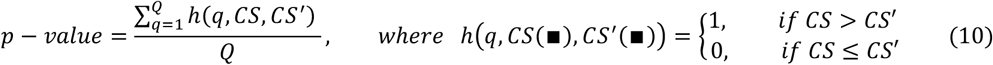

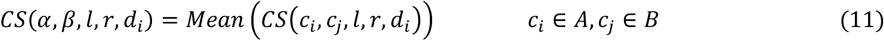

## III Results and discussion

### A. Experiment materials

Our computational experiments use the datasets published by Lofoff et al. (2022) which were generated by integrating seqFISH datasets from midline sagittal sections on three mouse embryos at the 8-12ss (corresponding to the embryonic day (E)8.5–8.75) with two existing scRNA-seq atlas [14]. The authors’ seqFISH dataset provided spatially resolved expressions of 387 target genes, but they used the comparable scRNA-seq atlas to impute missing genes of the spatially resolved cells and produced an extensive list of mouse embryo cell types. The integrated data was obtained from https://crukci.shinyapps.io/SpatialMouseAtlas/. All three integrated embryo datasets, namely, E1, E2 and E3, have 29452 genes and have 17806 cells, 14185 cells and 20577 cells, respectively. All these datasets are published with “best practice” total 23 cell types of cell labels as presented in Fig. 3. The figure also shows the number of cells included in each cell type along with the respective 2D plotting of cells with cell type colored accordingly. Noticeable is the distinctive shape of E2 which could be due to sectioning irregularity performed on their 2^nd^ embryo. Our rendering of this 2D cell type coloring confirms the authors’ suggestion that their 2^nd^ embryo suffers from depletion of mesodermal and tail-specific populations [14].

**Fig. 3.**
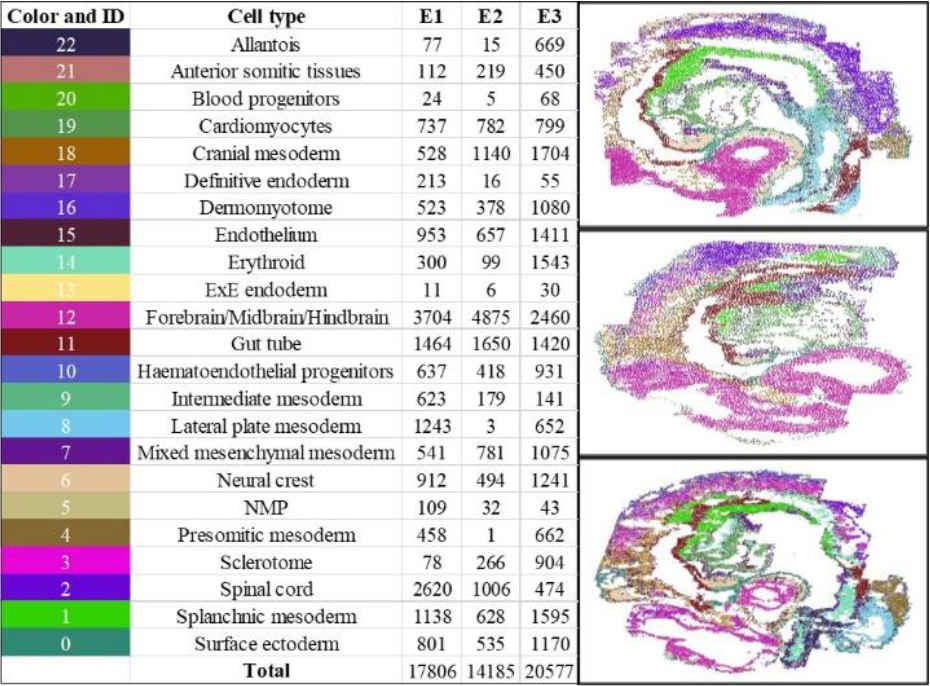
The details of E1, E2 and E3 datasets. Each datasets have 23 type cells.

### B. CGCom can effectively identify cell types

To assess the performance of the cell GAT classifier in CGCom, we use all three datasets as benchmarks. We explored the cell type prediction performance in two steps. First, we studied if and how the maximum communication ratio (*P*) may affect the classification accuracy. The grid search method is used with *P* ranging from 0.0001 to 0.02. The relationship between accuracy and *P* in all three datasets with 1% human-labeled samples are shown in Fig. 4. As shown, the GAT accuracy increases monotonically when *P* increases in the beginning, but after the knee of the curve (0.005) *P* no longer affects the classification accuracy. In all three datasets, the *P* with the best accuracy lands around 0.005 and we use this value as the best hyperparameter in our subsequent analyses. Noticeable in this performance analysis is that the plot for E2 stands out possibly due to block irregularity mentioned earlier. Nevertheless, the pattern of the knee of the curve does not change providing an added confidence in our way of choosing *P*.

**Fig. 4.**
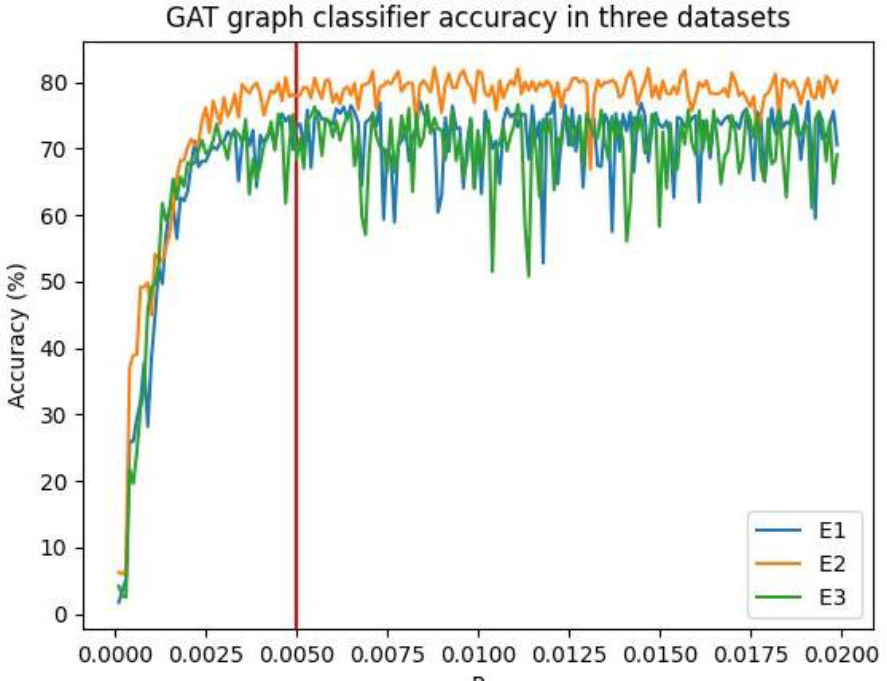
The relationship of accuracy and the max communication distance with 1%the training sample

Second, we conducted a comparative analysis by pitting our cell classifier against an MLP-based classifier using identical learning rates, the same number of hidden layers, and the same optimizer. Four criteria were employed to assess the performance of the classifiers including: Accuracy, Multi-class Precision, Multi-class Recall, and Multi-class F1-score. We varied the ratios of human-labeled samples at 1%, 5%, and 10%. As shown in Table 1, the performance of GAT and MLP increases when the ratio of human-labeled samples increases. Compared to MLP, GAT always outperformed MLP with ~30% improvement in testing Accuracy, and ~0.3 improvement in F1 score, Recall and Precision.

**Table 1.** The detailed performance comparing GAT with MLP for all three datasets under different training ration situation when P is set to 0.005.

| Training ratio | E1 | E2 | E3 |
| --- | --- | --- | --- |
|  | GAT |  |  |
| 1% | Accuracy: 77%, F1-score: 0.75, Precision: 0.75, Recall: 0.77 | Accuracy: 76%, F1-score: 0.77, Precision: 0.77, Recall: 0.79 | Accuracy: 69%, F1-score: 0.67, Precision: 0.69, Recall: 0.69 |
| 5% | Accuracy: 77%, F1-score: 0.76, Precision: 0.76, Recall: 0.77 | Accuracy: 79%, F1-score: 0.74, Precision: 0.78, Recall: 0.76 | Accuracy: 77%, F1-score: 0.76, Precision: 0.77, Recall: 0.77 |
| 10% | Accuracy: 82%, F1-score: 0.81, Precision: 0.82, Recall: 0.82 | Accuracy: 87%, F1-score: 0.87, Precision: 0.88, Recall: 0.87 | Accuracy: 80%, F1-score: 0.80, Precision: 0.80, Recall: 0.80 |
|  | MLP |  |  |
| 1% | Accuracy: 49%, F1-score: 0.42, Precision: 0.50, Recall: 0.49 | Accuracy: 55%, F1-score: 0.43, Precision: 0.37, Recall: 0.55 | Accuracy: 54%, F1-score: 0.46, Precision: 0.47, Recall: 0.54 |
| 5% | Accuracy: 54%, F1-score: 0.45, Precision: 0.42, Recall: 0.55 | Accuracy: 66%, F1-score: 0.58, Precision: 0.58, Recall: 0.67 | Accuracy: 56%, F1-score: 0.47, Precision: 0.50, Recall: 0.56 |
| 10% | Accuracy: 57%, F1-score: 0.54, Precision: 0.63, Recall: 0.58 | Accuracy: 66%, F1-score: 0.56, Precision: 0.49, Recall: 0.66 | Accuracy: 58%, F1-score: 0.50, Precision: 0.49, Recall: 0.59 |

Our conclusion is that CGCom’s outperformance over MLP is due to the former’s use extra information, that is, the cell type information of neighboring cells which is not used at all in the MLP model. The next question is then what if we instruct the model to pay a deeper “attention” to utilize ligand-receptor relationships embedded in gene expression values on top of the cell graph. Can the model now predict cell-cell communication patterns? This topic is further discussed in the following subsections.

### C. CGCom can identify cell communication

To assess the performance of CGCom in inferring cell communication, we first check communication coefficients between each pair of cell types. i.e., how frequently/infrequently cells of two types communicate with each other. Fig. 5a shows a summary view of such coefficients between a pair of three datasets, E1, E2 and E3, in the form of heatmaps. This heatmap rendering also exhibited the irregularity of E2, as we anticipated. Nevertheless, the analysis was performed on three comparable tissue blocks, their heatmap patterns should be consistent across all three cases and any such case would be considered more likely a true positive. Two cases are notable, one enclosed by small square blue boxes and one enclosed by small square red boxes, in Fig 5A. The first one is for communication coefficient between “Anterior somitic tissues” and “Intermediate mesoderm” which was high in all three cases. The related finding in the literature survey is “Somitic tissues can influence the development of intermediate mesoderm during embryogenesis” by Tani et. al. [15]. How this relationship is revealed in our spatial expression mapped image is shown in Fig. 5B1. The three expanded views enclosed by blue boxes show that somitic tissues cell are neighboring with intermediate mesoderm cells. The second pair for the high communication coefficient is between “Surface ectoderm” and “Anterior somitic tissues”. The related finding in the literature survey is “Surface ectoderm is necessary for the morphogenesis of somites” by Correia et. al. [16]. This relationship is also clearly visible in our spatial expression mapped image shown in Fig. 5B2. In all three expanded views enclosed by red boxes in Fig. 5B2, surface ectoderm cells are neighboring with anterior somitic tissues cells.

**Fig. 5.**
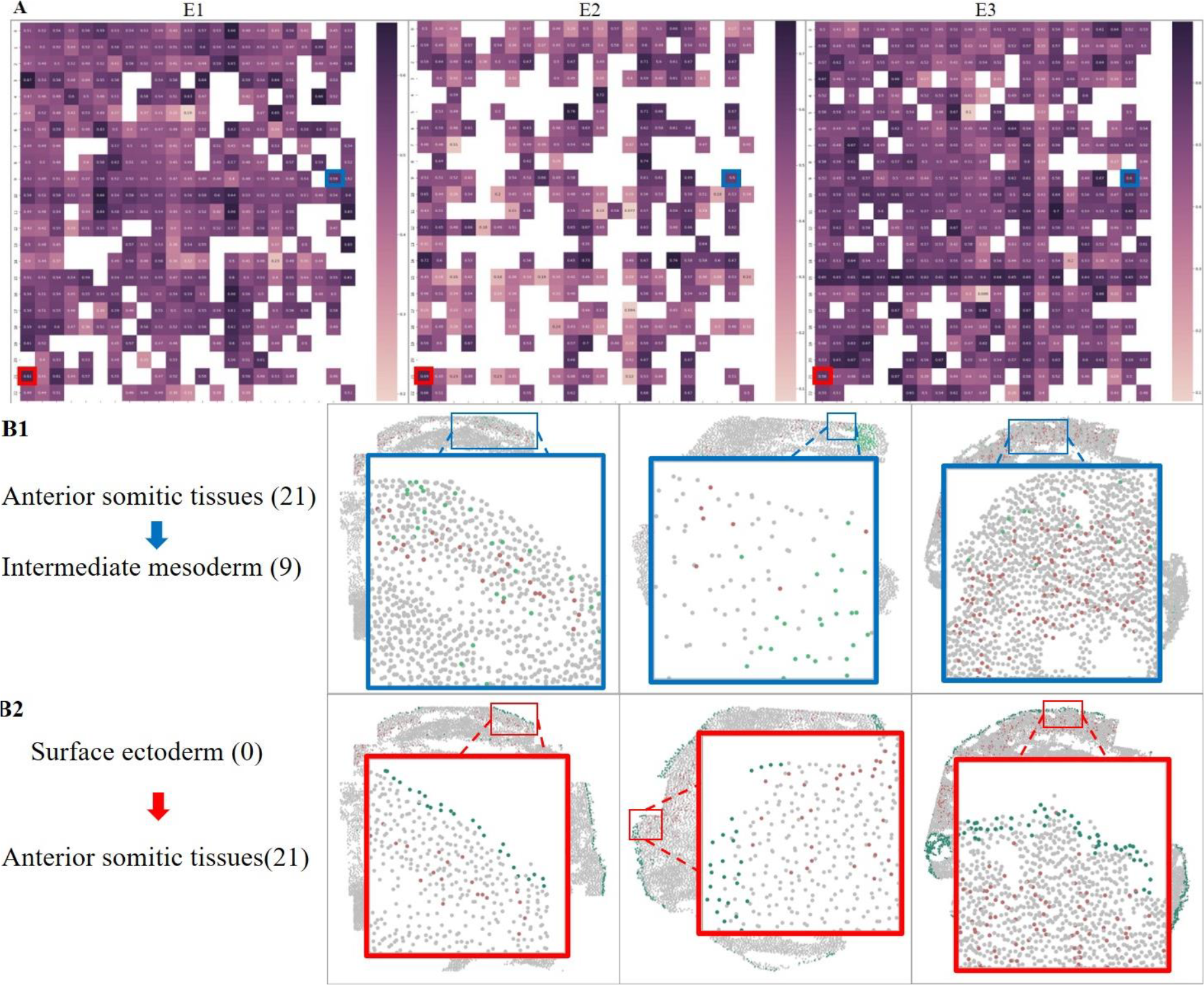
**A**. The attention score between each cell type pairs in all three datasets. **B**. The visualization of cell location in each dataset for the selected cell type pairs. The red box stands for the area that two types of cells are neighboring.

We also conducted a comparative analysis with two widely used methods, CellChat and CellPhoneDB. Fig. 6 shows this comparative study outcome in the form of Sankey diagrams as these can succinctly visualize the ligand and receptor communication patterns identified by all three datasets. In Fig. 6A1, each tile shows how ligands listed on the left interact with their corresponding receptors listed on the right. The bandwidth connecting two points in each side of the tile represents the communication score derived using each method. The same comparison analysis was performed on all three datasets, thus generating a total of 9 tiles. An interesting discovery in this comparison is that CGCOM and CellPhoneDB could be more immune to irregular sectioning (i.e., only E2 analysis outcome differs for CellChat). Examining differences in bandwidth also reveals which ligand and receptor pairs are statistically significant and more commonly appearing across different analyses. Common pairs appearing in all three datasets are individually shown in Fig. 6A2, 3 and 4 and they are enclosed by black boxes with genes names stated in them. These three subfigures also include two cases: one that is discovered only by CGCom, enclosed by green boxed (Fig 6A2); and one that is discovered by CellPhoneDB and CellChat, enclosed by red boxes (Fig 6A3 and Fig6A4, respectively). We discuss later that the one discovered by CellPhoneDB and CellChat is more likely a false positive. To further explore the communication patterns, Fig. 6B shows Venn diagrams visualizing how significantly the patterns obtained from the three methods overlap with each other. It shows that a total of 598 ligand-receptor pairs were commonly identified by all three methods. Regarding pair-wise comparisons, CellPhoneDB vs. Cellchat included a very large proportion patterns (2922) commonly but not so much between CellPhoneDB vs. CGCom and Cellchat vs. CGCom suggesting that CGCom is most discriminating (i.e, least overlapping with outcomes from other two methods).

**Fig. 6.**
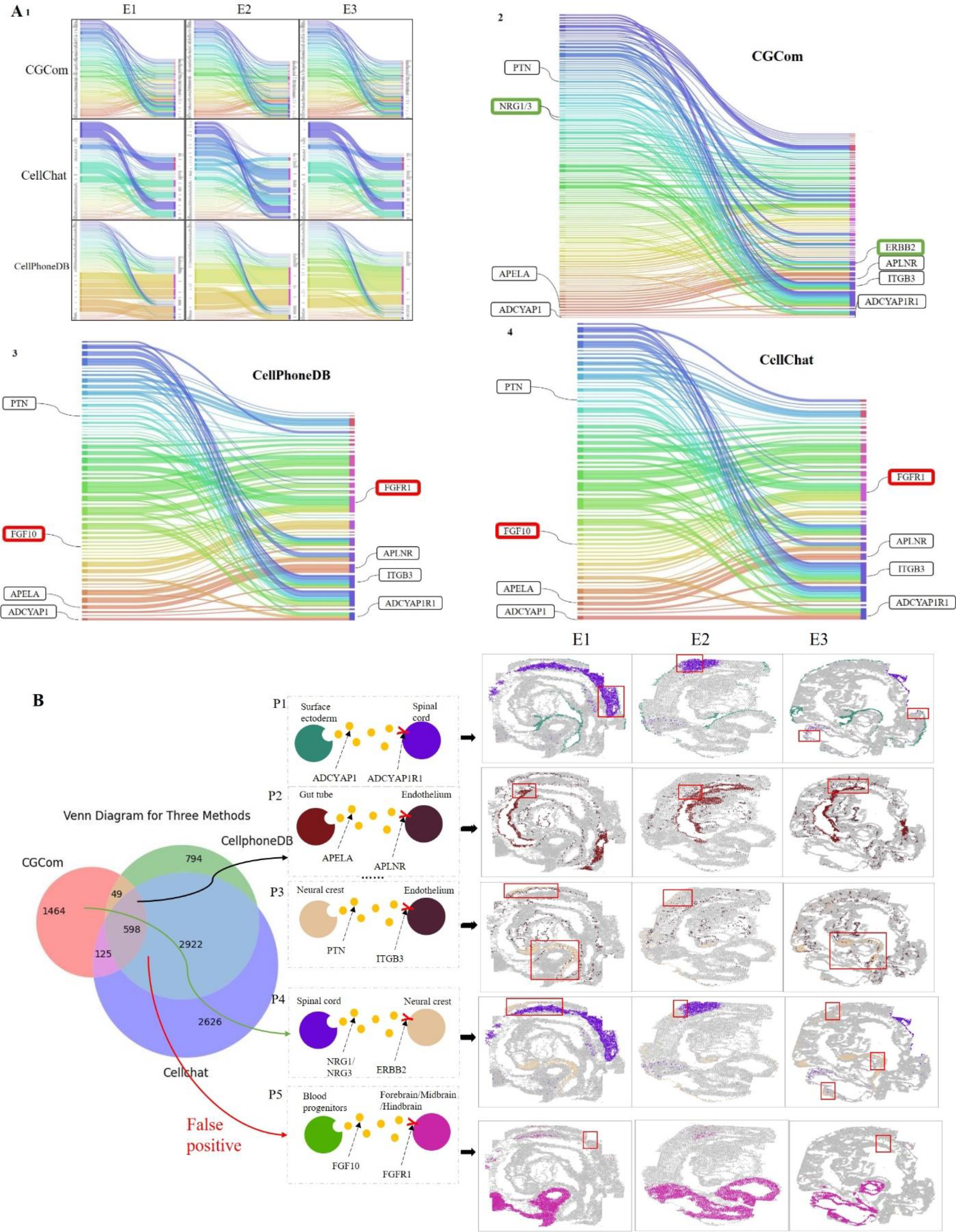
**A**. Sankey diagram for the ligand receptor using all three datasets with three different methods. **B**. The communication patterns details and visualization including three identified by all the methods.

Fig. 6B also shows five specific ligand-receptor communication pairs (P1 – P5) that we identified. Of these P1-P3 are predicted by all three methods (i.e., one of 598 common communication relationships). P1 involves “surface ectoderm cells” secreting ADCYAP1 (adenylate cyclase activating polypeptide1) ligand which is detected by ADCYP1R1 (ADCYAP receptor type 1) receptor. The work of Xu et al. in 2016 [17] states that ADCYAP1 plays a role in the proliferation of hematopoietic progenitor cells in murine bone marrow. Additionally, Mabuchi et al. in 2004 [18] reports ADCYAP’s importance in the development of spinal sensitization and neuropathic pain induction. Our results highlight that surface ectoderm cells may influence spinal cord cells through the ADCYAP1 signal. Fig 6B also includes where in our spatial expression mapped images for E1 – E3, this P1 relationship appears (enclosed by red rectangle boxes). P2 involves “gut tube cells” secreting the apelin receptor early endogenous ligand (APELA), whose signals are received by “endothelial cells” through the apelin receptor (APLNR). Helker et al. in 2020 [19] sates that Apelin signaling propels vascular endothelial cells into a pro-angiogenic state. The cells exhibiting P2 patterns are also captured in its corresponding spatial expression mapped images (enclosed by red rectangle boxes). P3 involves “neural crest cells” communicating with “endothelial cells” through pleiotrophin (PTN) and integrin subunit beta 3 (ITGB3). Heiss et al. [20] report that PTN may mediate the pro-angiogenic effects of endothelial progenitor cells.

Among the five patterns (P1 – P5), P4 is uniquely reported by CGCom. P4 involves “spinal cord” and “neural crest cells” through Neuregulin family gene (NRG1/NRG3) and erb-b2 receptor tyrosine kinase 2 (ERBB2). Piotrowski et. al. [21] states that spinal cord precursors utilize neural crest cell mechanisms to generate hybrid peripheral myelinating glia and one of the spinal cord cells (motor exit point glia) development depends on atonally derived NRG1. Despite this seemingly supporting relationship reported in the literature, P4 is one of 1464 relationships uniquely identified by CGCom.

Another noticeable pattern is P5 which is suggested between “blood progenitor cells” and brain cells through FGF10 and FGFR1. It belongs to one of 2922 relationships commonly discovered by CellChat and CellPhoneDB but not called by CGCom. The reason behind CGCom not calling this pattern is visible in our spatial expression mapped images for E1 and E3; here blood progenitor cells (enclosed by red boxes) are positioned very far away from Forebrain/Midbrain/ Hindbrain cells (pink colored cells). In both cases, these two cell types are not within the cell proximity used to train the model. In E2, only five blood progenitor cells were present. Is it truly a false positive relationship or not may require further study. Nevertheless, not calling P5 suggests that CGCom’s use of cell proximity information can discern proximity-driven communication pattern decision making.

Lastly, we discuss how the discovery of a relationship by CGCom can be explained in terms of gene expression statistics of interacting cell types. Fig. 7A illustrates the discovery of the PTN-ITGB3 communication relationship between “neural crest cells” and “endothelial cells” in E1. Its two large red boxes show expanded views of positional layout of cells (nodes), their types (by color, orange vs. black), and ligand-receptor pairing (edges between nodes). Shown in Fig 7B are the violin plots for gene expression levels of PTN and ITGB3 in both cell types (thus showing four panels of violon plots). Nural crest cells are high in PTN and ITBG3 are high in endothelial cells (gene expression pattern showing (above the red line indicating the mean z-score). The two genes can form a ligand-receptor pair is known along with their corresponding cell types encoded into edges interconnecting nodes. Inferring that the ligand signal from a neural crest cell is passed to the receptor of an endothelial cell is plausible and captured by the red arrow given in Fig 7B. Fig 7C boxplot illustrates another piece of information that is used in the inference. These boxplots show the distribution of pathway scores downstream of the ligand-bound receptor cells (orange colored) and the same for receptor signal weak/absent cells (blue colored), i.e., cells that may not receive ligand signals. It shows that all the ITGB3 downstream scores are higher than 0 (red line) indicating that the downstream pathway routes beyond the receptor could have been all activated. Unfortunately, receptor signal weak/absent cells (blue colored) also show activated downstream signals, suggesting limitations in utilizing the downstream pathway score for the inference because activation signals may originate from different sources—a topic which is notoriously difficult to unravel.

**Fig. 7.**
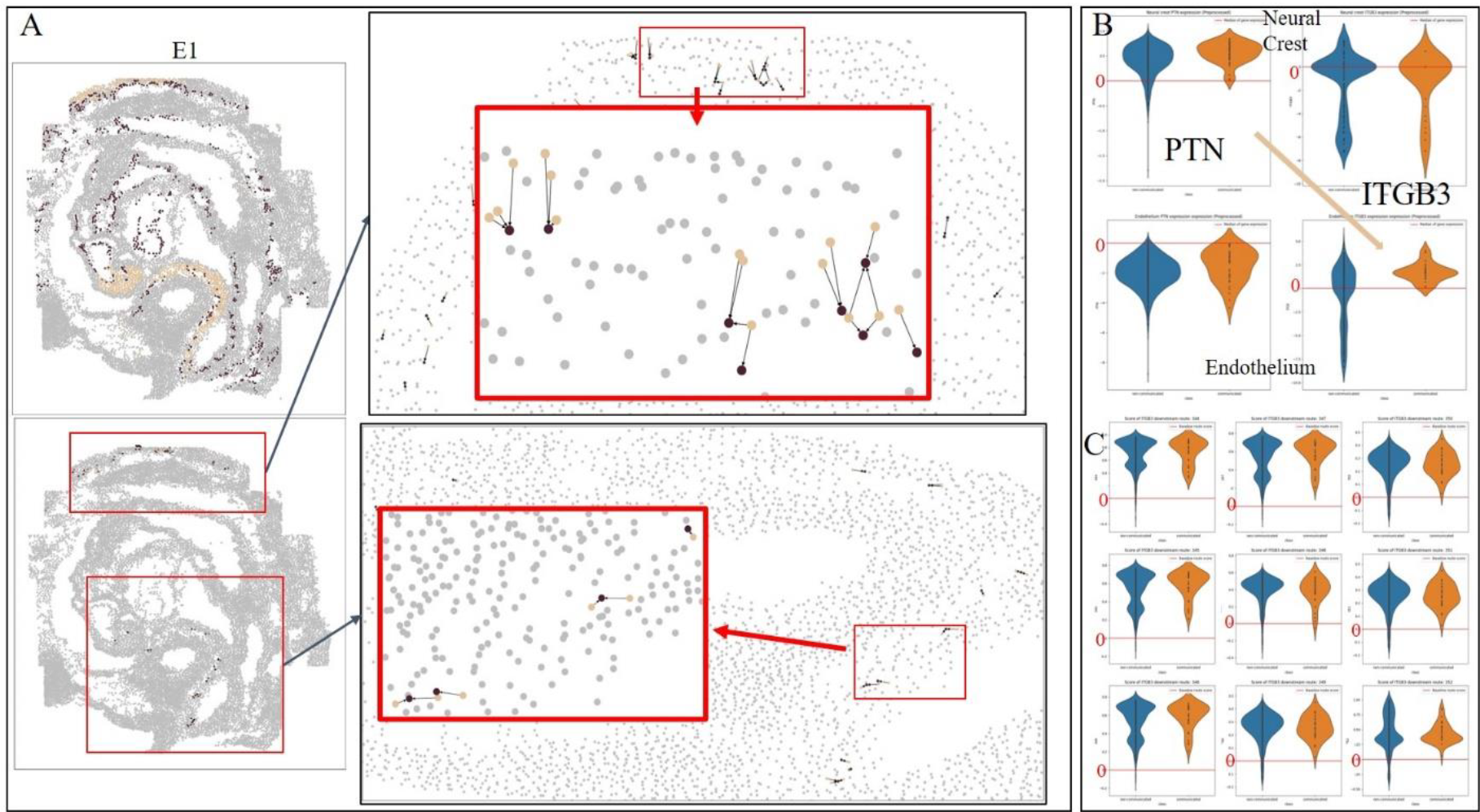
**A**. The visualization of paired neural crest and endothelium cell communication through PTN and ITGB3 in E1 dataset. **B**. The violin plot for the ligand and receptor expression value in Neural crest cell and endothelium. The orange violin stands for the communicating cell and blue violin stands for the non-communicating cell. **C**. 9 different downstream route scores of ITGB3 in the endothelium.

## IV conclusion and future work

Inferring cellular communication from spatial transcriptomics data is a challenging problem due to high-volume, high-sparsity, and noisy nature of data. CGCom is a deep learning method aiming to infer cell-cell communication by utilizing the two-dimensional x-y coordinate information available for each cell’s gene expression patterns. Specifically, CGCom exploits each cell’s ligand and receptor signal intensity information that can be combined with cell’s proximity information. By mapping the subcellular data from microscopic images to a graph, CGCom can take advantage of not only the cell’s proximity information but also the relationships between neighboring cells.

We reported a few benefits of CGCom. First, when comparing to the MLP model-based cell communication inference methods, CGCom provides a better performance in cell type classification. Second, CGCom integrates ligand and receptor relationship of neighboring cells into the learned model so that the attention scores at each layer can be translated into communication coefficients providing some degree of interpretability, that is, if nearby cells may or may not communicate with each other even through the ligand and receptor is highly expressed. Third, CGCom can accurately infer cell communication between not only pairs of individual cells but also pairs of cell types.

The current CGCom can be further extended. First, its edge weight scheme between a pair of neighboring cells can be extended beyond the current way of examining the ligand and receptor interaction relationships. For example, cells are known to intimately interact with the environment (i.e., extra cellular matrix) and the environment could fundamentally influence how cells communicate with each other. The current model is amenable to such an extension because incorporating data for the involved molecular elements (e.g., ECM deposited proteins, lipid molecules, etc.) can be easily done. Another direction for extension is enhancing the scoring of each cell’s state upon receiving signals from neighboring cells. Our current route-based pathway scoring to measure the activation/ suppression of receptor downstream activity is still quite limited. There are inherent difficulties in addressing this problem due to the sparse nature of data available from spatial transcriptomic assays. Nevertheless, pathway scoring methods such as rPAC [11] and even crosstalk inference tool such as ctBuilder [22] could offer ways to improve analyzing cellular states at the single cell level and thus enabling one to produce a better cell-cell communication prediction system.

## Code implement and availability

CGCom is implemented using Python 3.10 and Pytorch Geometric [23] and it will made open source and publicly available at GitHub (https://github.uconn.edu/how17003/CGCom) with the publication of this manuscript.

## Acknowledgment

Research reported in this work was supported in part by NIH Grant No. 4U54AR078664-03. Its contents are solely the responsibility of the authors and do not necessarily represent the official views of the NIH.

